# Early-life immune activation is a vulnerability factor for adult epileptogenesis in neurofibromatosis type 1

**DOI:** 10.1101/2023.10.31.564967

**Authors:** Rania Faidi, Aylin Y Reid

**Author notes:** **Correspondence:** Aylin Reid.

## Abstract

**Introduction:** Patients with Neurofibromatosis type 1 (NF1), the most common neurocutaneous disorder, can develop several neurological manifestations that include cognitive impairments and epilepsy over their lifetime. It is unclear why certain patients with NF1 develop these conditions while others do not. Early-life immune activation promotes later-life seizure susceptibility, neurocognitive impairments, and leads to spontaneous seizures in some animal models of neurodevelopmental disorders, but the central nervous system immune profile and the enduring consequences of early-life immune activation on the developmental trajectory of the brain in NF1 have not yet been explored. We tested the hypothesis that early-life immune activation promotes the development of spatial memory impairments and epileptogenesis in a mouse model of NF1.

**Methods:** Male wild-type (WT) and *Nf1*^*+/-*^ mice received systemic lipopolysaccharide (LPS) or saline at post-natal day 10 and were assessed in adulthood for learning and memory deficits in the Barnes maze and underwent EEG recordings to look for spontaneous epileptiform abnormalities and susceptibility to challenge with pentylenetetrazole (PTZ).

**Results:** Whereas early-life immune activation by a single injection of LPS acutely elicited a comparable brain cytokine signature in WT and *Nf1*^*+/*-^ mice, it promoted spontaneous seizure activity in adulthood only in the *Nf1*^*+/*-^ mice. Early-life immune activation affected susceptibility to PTZ-induced seizures similarly in both WT and *Nf1*^*+/-*^mice. There was no effect on spatial learning and memory regardless of mouse genotype.

**Discussion:** Our findings suggest *second-hit* environmental events such as early-life immune activation may promote epileptogenesis in the *Nf1*^*+/-*^ mouse and may be a risk-factor for NF1-associated epilepsy.

## 1 Introduction

Neurofibromatosis type 1 (NF1) is a neurocutaneous disorder caused by autosomal dominant germline mutations to the *NF1* gene (Viskochil et al., 1990; Wallace et al., 1990), a tumor suppressor gene on chromosome 17 encoding the protein neurofibromin. Neurofibromin is a multi-domain protein that is best characterized for its role as a Ras-guanosine triphosphate activation protein (Ras-GAP) (Weiss et al., 1999). NF1-associated pathogenic mutations lead to reduced or complete absence of neurofibromin, promoting constitutive activation of GTP-Ras and overactivation of cell proliferation and survival pathways. NF1 is a progressive condition and patients often exhibit a wide range of dermatological, cardiovascular, and musculoskeletal comorbidities which present gradually over their lifetime, severely affecting their quality of life. Pigmentary lesions such as neurofibromas and café au lait macules (CALMs) are the most common clinical manifestations of the disease (Shofty et al., 2015). Less understood are the neurological manifestations of NF1, which include cognitive impairments and epilepsy.

Epilepsy occurs at a higher incidence in NF1 patients (5-11%) (Bernardo et al., 2020; Sorrentino et al., 2021; Hébert et al, 2022) than in the general population (1-2%) (Hauser, 1995). Whereas structural abnormalities have been associated with seizures in some cases (Hsieh et al., 2011; Pecoraro et al., 2017; Bernardo et al., 2020), they do not account for the manifestation of seizures in *all* NF1-associated epilepsy cases. For instance, only 50% of reported NF1-associated epilepsy cases are structural epilepsies resulting from brain abnormalities detectable on neuroimaging (Bernardo et al., 2020) and structural NF1-associated lesions on MRI do not always co-localize with epileptiform discharges on EEG (Serdaroglu et al., 2019). In fact, many reports of NF1-associated epilepsy lack detailed electroencephalographic (EEG), clinical, and brain imaging correlates that would confirm the epileptogenicity of a structural lesion (Bernardo et al., 2020; Kulkantrakorn and Geller, 1998). This underscores an uncertainty around whether an underlying structural lesion is always the *true* epileptogenic basis of NF1 epilepsy, or whether there are molecular alterations, driven by disease-associated loss-of-function mutations in *Nf1*, that mediate NF1-associated hyperexcitability. Until very recently, research on seizure susceptibility and epileptogenesis in NF1 was lacking. Recent work from our group with the non-lesional *Nf1*^*+/-*^ mouse model demonstrated increased seizure susceptibility and increased rates of epileptogenesis (Rizwan et al, 2019; Sabetghadam et al., 2020).

In addition to epilepsy, cognitive impairments are also a common neurological manifestation of NF1, occurring in up to 65% of patients (Rosser et al., 2003) and including learning disabilities, attention deficits, and social perception problems (Hyman et al., 2005). Animal models of NF1, including the non-lesional *Nf1*^*+/-*^ mouse, replicate these learning impairments in attention and spatial memory tests (Silva et al., 1997; Li et al, 2005), and have been shown to be mediated by RAS-hyperactivity downstream of neurofibromin, which promotes alterations in GABAergic and dopaminergic function (Costa et al., 2002; Cui et al., 2008; Diggs-Andrews et al., 2013). It is unclear why some patients with NF1 go on to develop cognitive impairments and epilepsy, while others do not.

There is an abundance of clinical and experimental evidence suggesting neuroinflammation is involved in the pathogenesis of seizures and may promote cognitive impairments in learning and memory (Ravizza et al., 2018; Maroso et al., 2010). In particular, epidemiological studies have suggested exposure of the developing fetus to immune challenges sustained in the mother during early to mid-pregnancy result in the greatest risk in the offspring for epilepsy (Sun et al., 2008; 2011) and neurodevelopmental disorders, such as autism and schizophrenia (Solek et al., 2018; Guma et al., 2019). Similarly, early-life *in vivo* studies suggest that activating the innate immune system during a critical window between postnatal day (P)7-14 (equivalent to early human infancy) with systemic lipopolysaccharide (LPS) is associated with an increased later-life susceptibility to chemo-convulsant induced seizure activity (Galic et al., 2008; Harré et al., 2008; Chen et al., 2013) and altered expression of NMDA receptor subunits (Harré et al., 2008), with mixed effects on hippocampal dependent memory (Chen et al., 2013; Dinel et al., 2014). Early-life systemic immune activation in a mouse model of autism spectrum disorder (ASD) has also been shown to promote spontaneous seizure activity in adulthood (Lewis et al., 2018).

There are reports of an increased level of peripheral inflammation in patients with NF1 (Farschtschi et al., 2016; Torres et al., 2016; Park et al., 2013; Lasater et al., 2010). It is therefore intriguing to determine if immune activation in NF1, particularly early in life, is associated with an increased susceptibility to cognitive impairments and seizures. In the present study we used the non-lesional *Nf1*^*+/-*^ mouse model to determine whether early-life immune activation exacerbates the seizure susceptibility and spatial memory impairments associated with NF1.

## 2 Material and Methods

Experiments were performed in accordance with the policies and guidelines established by the Canadian Council on Animal Care and ethical approval from the University Health Network-Animal Resources Centre. *Nf1*^*+/-*^ mice (strain #008192) were originally purchased from Jackson Laboratory (Bar Harbor, ME, USA), and bred in house. *Nf1*^*+/+*^ littermates were designated as the WT controls and are henceforth referred to as such. Only male offspring mice were used in this study. The colony was maintained under standard specific pathogen-free conditions at 20-21°C on a 12h light/dark cycle with food and water available *ad libitum*. To mitigate potential litter effects, 14 litters from a total of 13 different breeding pairs contributed pups to this study. An overview of the experimental design is presented in Figure 1.

**Figure 1.**
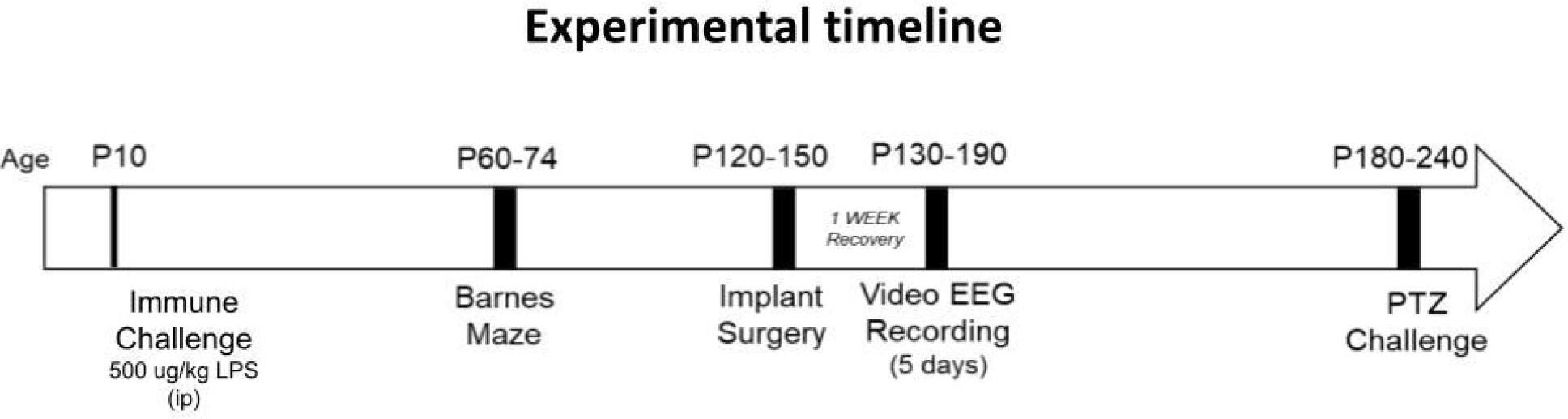
Experimental Timeline. Postnatal day (p)10 mice sacrificed four hours following LPS challenge, for cytokine analysis on brain tissue. Some mice were allowed to mature after p10, for assessment of later-life spatial memory (between p60-p74), spontaneous recurrent seizure activity (p130-p190) and induced seizure activity (p180-p240).

### 2.1 Postnatal Lipopolysaccharide Challenge

P10 *Nf1*^*+/-*^ and WT littermate male pups received a single intraperitoneal injection of 500 ug/kg lipopolysaccharide (Escherichia coli, serotype O26:B6, Sigma-Aldrich, Oakville, Canada) or a matched volume of 0.9% saline solution (sterile vehicle). Whenever possible, both *Nf1*^*+/-*^ and WT males were used from each litter. Whereas a milder dose of 100 ug/kg LPS has been reported to induce early-life immune activation in rodents (Galic et al., 2008; Lewis et al., 2018), our own preliminary experiments with this dose did not reveal any changes in brain cytokine levels of LPS-challenged versus saline-treated WT and *Nf1*^*+/-*^ controls. Therefore, to induce immune activation, the dose of LPS was increased to 500 ug/kg (100 ug/mL in sterile normal saline). Immediately following injection, all mice were returned to their dams with cages places on heating pads. Saline control and immune-challenged pups were included in each litter to minimize inter-litter variation in maternal care. A subset of mice were sacrificed four hours after saline or LPS injection for analysis of brain pro-inflammatory cytokine levels. The other subset was used for behavioral studies and EEG recordings in adulthood. These mice were weaned at P21 and housed with up to five same-sex littermates with different genotype and postnatal treatment combinations, at which point the experimenter remained blind to the experimental groups.

### 2.2 Neuroinflammatory Cytokine Profiling

Four hours following the immune challenge, whole-brain tissue (excluding cerebellum and olfactory accessories) was extracted on ice from P10 pups (n=5 per group), flash frozen, and stored at -80°°C. Brain tissue was mechanically homogenized by a cordless motor pellet mixer (#V8185-904; VWR, Mississauga, Canada) in a PBS-based cell lysis buffer complete with EDTA-free protease inhibitors (#04693159001; Roche, Mississauga, Canada) and 1 mmol/L phenylmethylsulfonyl fluoride (#PMSF123.5; BioShop, Toronto, Canada). Tissue lysates were analyzed for the concentration of total protein and assayed in a Mouse Magnetic Luminex Screening Assay (LXSAMSM, R&DSYSTEMS, Oakville, Canada) for the concentration of the following cytokines: TNF-α, IL-1β, IFN-γ, and IL-6. Refer to Appendix S1 for supplementary methods.

### 2.3 Barnes Maze

Between the P60-74 window, adolescent control and immune-challenged mice (n= 8-11 per group) were tested in the Barnes Maze to assess spatial learning and memory, as previously described (Sunyer et al., 2007). Mice were transported to the testing room 24 hours before the first training day, and remained in their littermate cages, in order to habituate to the novel holding conditions. Mice were spatially trained for a total of 16 training trials over four days (acquisition phase), and short-term reference memory was tested once at 24 hours following spatial acquisition. All groups were trained and tested during the light phase, and the maze was cleaned with 70% isopropyl alcohol between trials to prevent the transfer of olfactory cues. Smart TriWise video tracking system (Version 3.0.06, Panlab Harvard Apparatus, Massachusetts, USA) was used to record all trials and acquire the data. Refer to Appendix S1 for supplementary methods outlining the experimental protocol and the acquired parameters.

### 2.4 Video-EEG monitoring

Mice were maintained in the vivarium after Barnes maze testing until a window between P120-150 for video-EEG monitoring, at which point five mice per group underwent implantation of intracranial electrodes. Briefly, mice were anesthetized with 2% isoflurane and fixed to a stereotaxic frame. The skull was exposed through a skin incision and small burr holes were drilled to implant a set of bipolar polyamide-insulated twisted stainless-steel wire electrodes (outer diameter 150 um; Plastics One, Roanoke, USA) in the CA3 region of the hippocampus (bregma: 2.5mm posterior, 3.0 mm lateral, 3.0 mm depth) and the contralateral sensorimotor neocortex (2.0 mm posterior, 2.0 mm lateral, 1.5 mm depth). One reference electrode was implanted at a frontal area (1.0 mm anterior, 1.0 mm lateral, 0.5 mm depth). Following surgery, mice were allowed to recover for one week before video-EEG monitoring commenced, as previously described (Rizwan et al., 2019).

Video EEG-monitoring was conducted between P130-190. The implanted mice were placed individually in modified cages with food and water *ad libitium*. A slip-ring commutator was mounted on top of the cage and connected to the mouse through a flexible cable which was of sufficient length to allow the mouse to move freely throughout the cage. The EEG signal was sampled by one of two sets of MP160 amplifiers (BIOPAC Systems, Inc., California, USA) with a sampling rate of either 250 Hz, bandpass filtered with a high-pass of 1.0Hz and low-pass of 3.0K Hz (MP160-194D) or a sampling rate of 1000Hz, bandpass filtered with a high-pass of 0.5Hz and low-pass of 100Hz (MP160-1A88). A webcam (LINKIT security web cameras; BIOPAC Systems, Inc., California USA) was anchored next to each cage to continuously monitor the animal ‘s behavior. Video-recordings were time-stamped to synchronize with the EEG recordings, and were acquired, stored and analyzed using the AcqKnowledge software (Version 5.0; BIOPAC Systems, Inc., California USA). Continuous recordings were carried out 24 hours daily for 5 consecutive days. A two-hour epoch from each of the light and dark cycles was visually inspected for interictal spikes. Entire recordings were analyzed for spontaneous recurrent seizure activity. EEG analyses were confirmed by two independent reviewers to minimize bias. An interictal spike was defined as a highly contoured waveform that appeared at least twice the baseline amplitude and lasted between 20-70 ms. A seizure was defined as rhythmic discharge events with an amplitude greater than four times the baseline and lasting a minimum of five seconds. Any ictal spikes (occurring as part of a seizure) were not included in the separate interictal spike counts.

### 2.5 Pentylenetetrazol-induced seizure susceptibility

Between the P180-240 window, seizure susceptibility to the GABA antagonist pentylenetetrazol (PTZ) was assessed (n=6-8 per group). In order to determine a mildly convulsive dose of PTZ that would enable us to differentiate the seizure susceptibility between our groups, a number of doses of PTZ were tested on naïve (i.e. have not undergone early-life immune challenge and were not implanted with intracranial electrodes) *Nf1*^*+/-*^ and WT animals, and in a subset of experimental *Nf1*^*+/-*^ and WT animals. Doses of 30, 40, 50 and 60 mg/kg of PTZ were tested. Based on preliminary findings of no seizures at lower doses, and status epilepticus followed by death at higher doses, a dose of 40 mg/kg of PTZ (i.p.; P6500-100G, Sigma-Aldrich, Oakville, Canada) was selected as a sub-convulsive dose, and mice were challenged as previously described (Van Erum et al., 2019). Immediately following injection, every animal was monitored for one hour and video-EEG recordings were acquired and visually inspected as described above. The latency to the first spike, the first epileptiform discharge event (EDE), and the first induced-seizure were determined. Spikes and seizures were defined as described above, EDEs were defined as high amplitude rhythmic discharges containing bursts of slow wave, spike-wave and/or polyspike wave discharges and lasting less than five seconds. Additionally, the total count of spikes, total count of EDEs, and cumulative induced-seizure duration during the period of monitoring were determined.

### 2.6 Statistical Analysis

For all experiments, data are reported as mean ± standard error of the mean (SEM). GraphPad Prism 8 statistical software (Version 8.4.3, California, USA) was used for statistical analyses. Cytokine concentrations were analyzed in a two-way analysis of variance (ANOVA). Barnes Maze acquisition data were analyzed using a two-way repeated measures ANOVA. The assumption of equal variances (sphericity) was not assumed, and the data were corrected by the Greenhouse-Geisser correction. Barnes maze probe data, spontaneous epileptiform abnormalities, and PTZ susceptibility data were compared in a two-way ANOVA. The proportion of animals that did/did not develop a seizure in response to PTZ were compared with a Chi-Square test. *p<*0.05 was considered significant for all analyses.

## 3 Results

### 3.1 Postnatal LPS increases levels of brain TNF-α and IL-6 in Nf1^+/-^ and WT mice

Analysis of brain tissue harvested four hours after injection of saline or LPS revealed no significant differences in brain cytokine levels between WT+Saline and *Nf1*^*+/-*^+Saline mice [IL-1β (F_(1,16)_=0.17, *p*=0.685); TNF-α (F_(1,16)_=0.45, *p*=0.510); IL-6 (F_(1,16)_=0.03, *p*=0.863) (Figure 2). Postnatal LPS induced a robust expression of IL-6 (F_(1,16)_=24.65, *p*<0.0001) and TNF-α (F_(1,16)_=21.70, *p*<0.001) in both *Nf1*^*+/-*^ and WT mice. Brain IL-1β levels were significantly elevated only in *Nf1*^*+/-*^+LPS mice (p=0.031), remaining comparable in WT+LPS *vs*. WT+Saline mice (*p*=0.193). No between-group differences or any significant treatment x genotype interaction effects for any of the assessed cytokines were observed.

**Figure 2.**
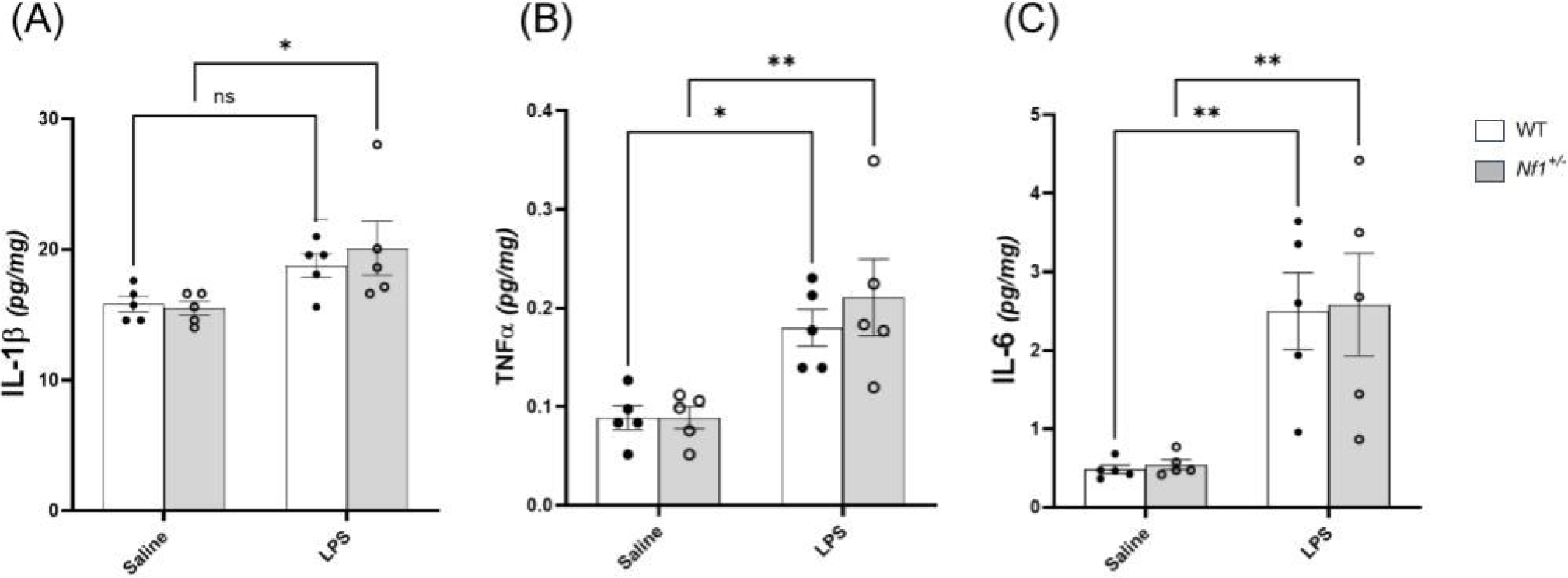
Postnatal LPS increases levels of brain TNF-α and IL-6 in *Nf1*^*+/-*^ and WT mice. There was a significant increase in the concentration of the pro-inflammatory cytokines IL-1β (A), TNF-α (B), IL-6 (C) in whole-brain tissue collected four hours after LPS challenge in p10 *Nf1*^*+/-*^ mice as compared to saline-treated controls. WT+LPS mice had similar increases in TNF-α (B) and IL-6 (C) compared wot WT+Saline controls, but no significant increase in IL-1β concentration (A). No significant interaction effect of treatment x genotype for any cytokine was observed. Data represent mean concentrations (pg/mg) ± SEM. \**p* < 0.05 and \*\**p* < 0.01 compared to saline controls.

### 3.2 Early-life immune activation did not affect spatial learning

All groups demonstrated ability to learn the Barnes maze over a four day training period. WT+Saline and *Nf1*^*+/-*^+Saline mice had significantly shorter primary latencies (WT *p*=0.003; *Nf1*^*+/-*^ *p*=0.001), shorter distances to the target (WT *p*=0.002; *Nf1*^*+/-*^ *p*=0.009) and made fewer primary errors (WT *p*=0.020; *Nf1*^*+/-*^ *p*=0.018) by Day 4 *vs*. Day 1 of training in the Barnes maze (Figure 3). Similarly, WT+LPS and *Nf1*^*+/-*^+LPS mice also exhibited shorter primary latencies (WT *p*<0.001; *Nf1*^*+/-*^ *p*=0.001), fewer primary errors (WT *p*=0.01; *Nf1*^*+/-*^ *p*=0.005) and shorter distance to the target (WT *p*=0.002; *Nf1*^*+/-*^ *p*=0.002) by Day 4 *vs*. Day 1 of training (Figure 3). Lastly, all groups acquired the maze at a similar rate, regardless of genotype or treatment, considering that no significant between-group differences on *any day of training* in primary latency (Figure 3, A.), nor in distance to the target (Figure 3, B.) were noted. Still, *Nf1*^*+/-*^+LPS mice were observed to have made significantly more primary errors (Figure 3, C.) than WT+LPS mice on Day 2 of training (*p*=0.022).

**Figure 3.**
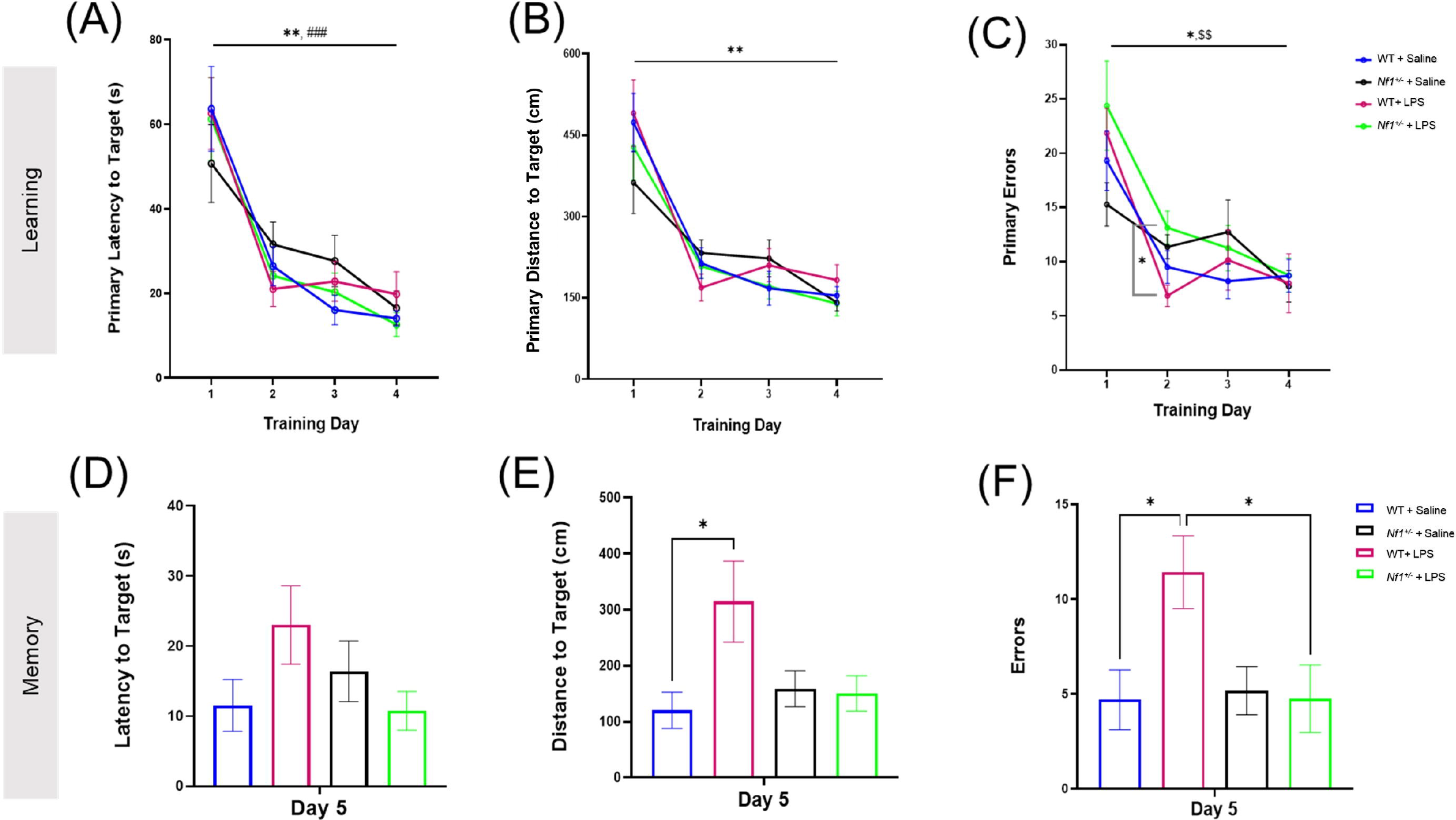
Early life inflammation does not affect spatial learning and memory in adult *Nf1*^*+/-*^ mice. The WT+Saline, *Nf1*^*+/-*^+Saline, WT+LPS, and *Nf1*^*+/-*^+LPS groups all learned the Barnes maze as demonstrated by improvements in primary latency (s) to the target hole (A), primary distance travelled (cm) to the target hole (B), and number of primary errors (C) over the four training days. The training phase was followed by a probe trial to assess short-term spatial memory on Day 5. There were no differences in the latency to the target hole between any groups (D). The WT+LPS group travelled a significantly longer distance to reach the target hole (E) and made more errors (F) than the WT+Saline controls. Data are reported as the mean ± SEM. [(*) was used to indicate that all groups were significantly different on Day 4 vs. Day 1: *p < 0.05, **p< 0.01, and ***p< 0.001 #p<0 .05. Specific within-group differences were represented with the following symbols: (#) for WT+Saline, ($) for *Nf1*^*+/-*^+Saline, (%) for *Nf1*^*+/-*^+LPS].

### 3.3 Early-life immune activation affected spatial memory in adult WT but not Nf1^+/-^ mice

Short-term spatial reference memory was assessed 24 hours after the last training trial (Day 5). There was no effect of genotype, and both WT+Saline and *Nf1*^*+/-*^+Saline mice demonstrated comparable short-term spatial memory evidenced by similar latency, distance, and total errors to the target hole (Figure 3, D-F). There was an effect of early-life immune challenge in WT mice only, as WT+LPS mice traveled longer distances (*p*=0.012) and made significantly more errors (*p*=0.030) than WT+Saline mice. A similar effect of LPS was not seen in *Nf1*^*+/-*^ mice. No between-group differences were observed in the latency to the target hole across all groups (Figure 3, D). The time spent exploring the Target quadrant during the Barnes maze probe trial was also analyzed to capture other potential differences in performance. All groups demonstrated a degree of bias towards exploring the target quadrant over other maze quadrants, and the percentage of time mice spent exploring the Target quadrant was comparable across all groups regardless of genotype or treatment (Appendix S2, Supplementary Figure 1.).

### 3.4 Early-life immune activation increased the number of spontaneous epileptiform spikes and promoted spontaneous seizure development in adult Nf1^+/-^ but not WT mice

The number of spontaneous interictal epileptiform spikes was measured in epochs of EEG recordings including both light and dark cycles. There was a significant effect of genotype with a greater number of average spikes in *Nf1*^*+/-*^ mice (*p*=0.027), and a significant effect of early-life immune activation with a greater number of average spikes in LPS-treated mice (*p*=0.0058). There was also a significant interaction (*p*=0.0084). Multiple comparisons showed no effect of LPS in WT mice (p=0.98), whereas treatment with LPS significantly increased average spike number in *Nf1*^*+/-*^ mice (*p*=0.0019). The proportion of interictal spikes in the light and dark phases was comparable (Figure 4, A.).

**Figure 4.**
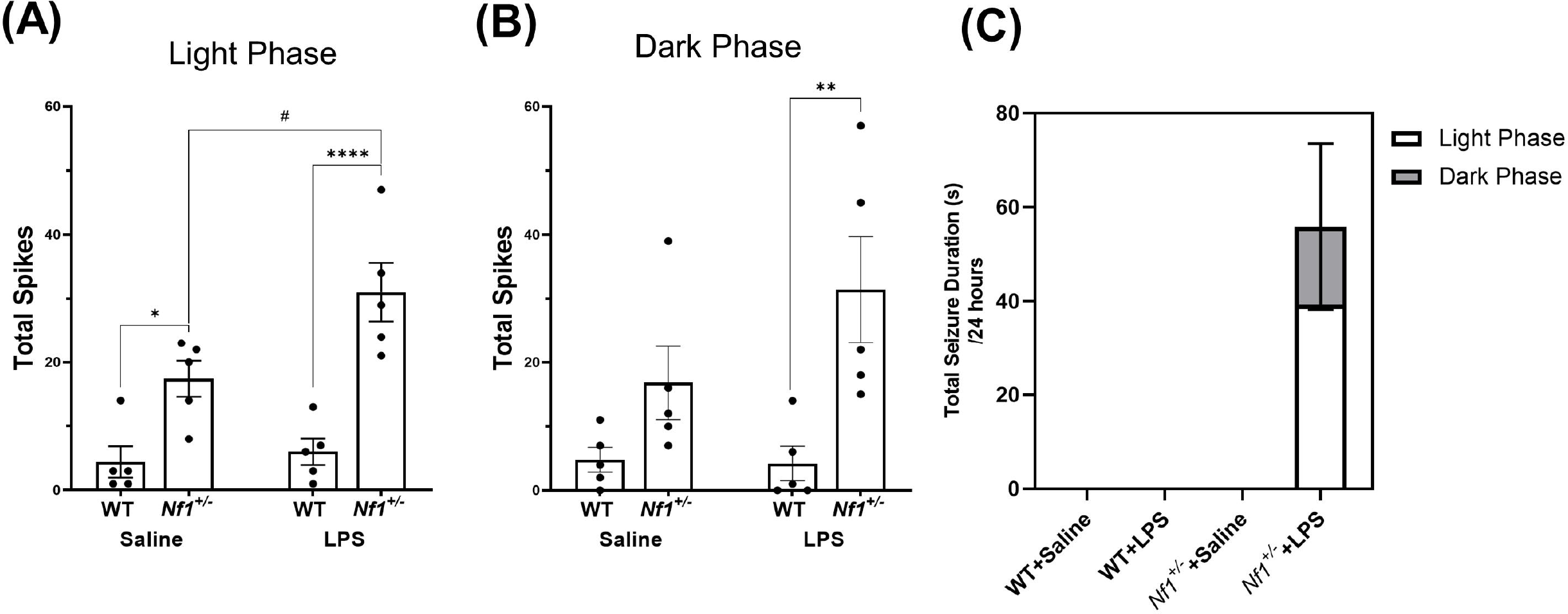
Early-life immune activation increases spike count and promotes spontaneous seizure activity in *Nf1*^*+/-*^ mice. A two-hour epoch of EEG recording was analyzed during the light (10:00-12:00) (A) and dark cycle (22:00-24:00) (B) for each animal. The *Nf1*^*+/-*^+Saline group had significantly more interictal epileptiform spikes during the light cycle than the WT+Saline group (A). The *Nf1*^*+/-*^+LPS group had more spikes than the WT+LPS group in both the light and dark cycles (B). Spontaneous seizures were only detected in the *Nf1*^*+/-*^+LPS group (C). Data are presented as mean ± SEM. *p < 0.05, **p< 0.01, and ****p< 0.0001 compared to another genotype group, #p<0 .05 compared to another treatment group.

Spontaneous seizure activity was not observed at any time in WT+Saline, WT+LPS, or *Nf1*^*+/*^+Saline mice. Spontaneous seizure activity was, however, observed in four out of five *Nf1*^*+/*^+LPS mice (Figure 4, B). There was a significant effect of genotype with a greater average seizure duration in *Nf1*^*+/-*^ mice (*p*=0.0219), and a significant effect of early-life immune activation with a greater average seizure duration in LPS-treated mice (*p*=0.0219). There was also a significant interaction (*p*=0.0219). Multiple comparisons showed *Nf1*^*+/-*^+LPS mice had a greater average seizure duration than *Nf1*^*+/*^+Saline mice (*p*=0.0077) or WT+LPS mice (*p*=0.0020). A greater proportion of seizures occurred during the light phase than the dark phase.

### 3.5 Early-life immune activation decreased the latency to the first spike after PTZ challenge

To assess hyperexcitability, we evaluated the proportion of animals that developed seizures in response to the PTZ challenge, as well as the latency to the first seizure, the cumulative seizure duration, the latency to the first spike, total spike count, the latency to the first EDE, and total EDE count in each group. An example of the different epileptiform events that were assessed is shown in Figure 5. As we used a subconvulsive dose of PTZ, not all animals developed electrographic seizures following PTZ challenge, and animals that did develop seizures showed varying behavioral severity. Two mice (one *Nf1*^*+/-*^+LPS and one WT+Saline) developed very severe seizures following PTZ-challenge and died from status epilepticus during testing. These were excluded from the analysis of the total number of spikes and EDEs but were included in the analysis of the latencies of these measures and any seizure-related parameters.

**Figure 5.**
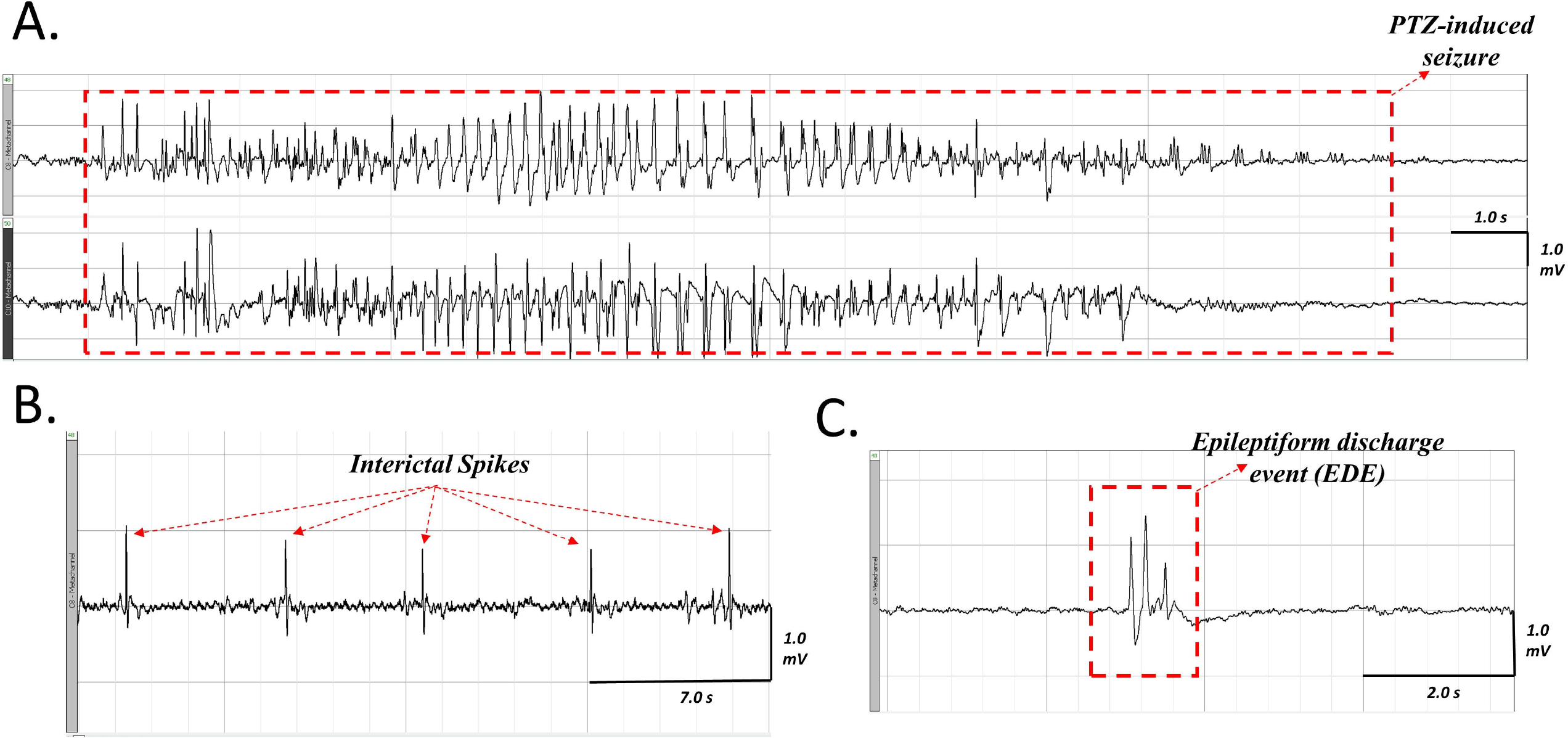
Representative ictal and interictal epileptiform activity observed on EEG following PTZ-challenge in an LPS-challenged *Nf1*^*+/-*^ mouse. A seizure occurring after PTZ challenge (A), the top tracing is the neocortical electrode and the bottom tracing is the hippocampal electrode. Interictal spikes (B) appear as sharp, contoured, and fast waveforms that are a minimum twice the baseline amplitude. An epileptiform discharge event (C) appears as a complex waveform (e.g. a polyspike wave discharge) that lasts less than five seconds.

There was a significant effect of early-life immune activation on the latency to the first spike, with a lower latency in mice receiving LPS (*p*=0.001), but no effect of genotype (*p*=0.2482) and no interaction (*p*=0.6042). Both WT+LPS (*p*=0.0271) and *Nf1*^*+/-*^+LPS (*p*=0.0151) groups had lower latencies to first spike than their respective saline controls. No significant differences were found in the total spike count across all groups (*p* = 0.791) (Figure 6, B.), or in the latency to the first EDE or total EDE count (Appendix S2, Supplementary Figure 2.).

**Figure 6.**
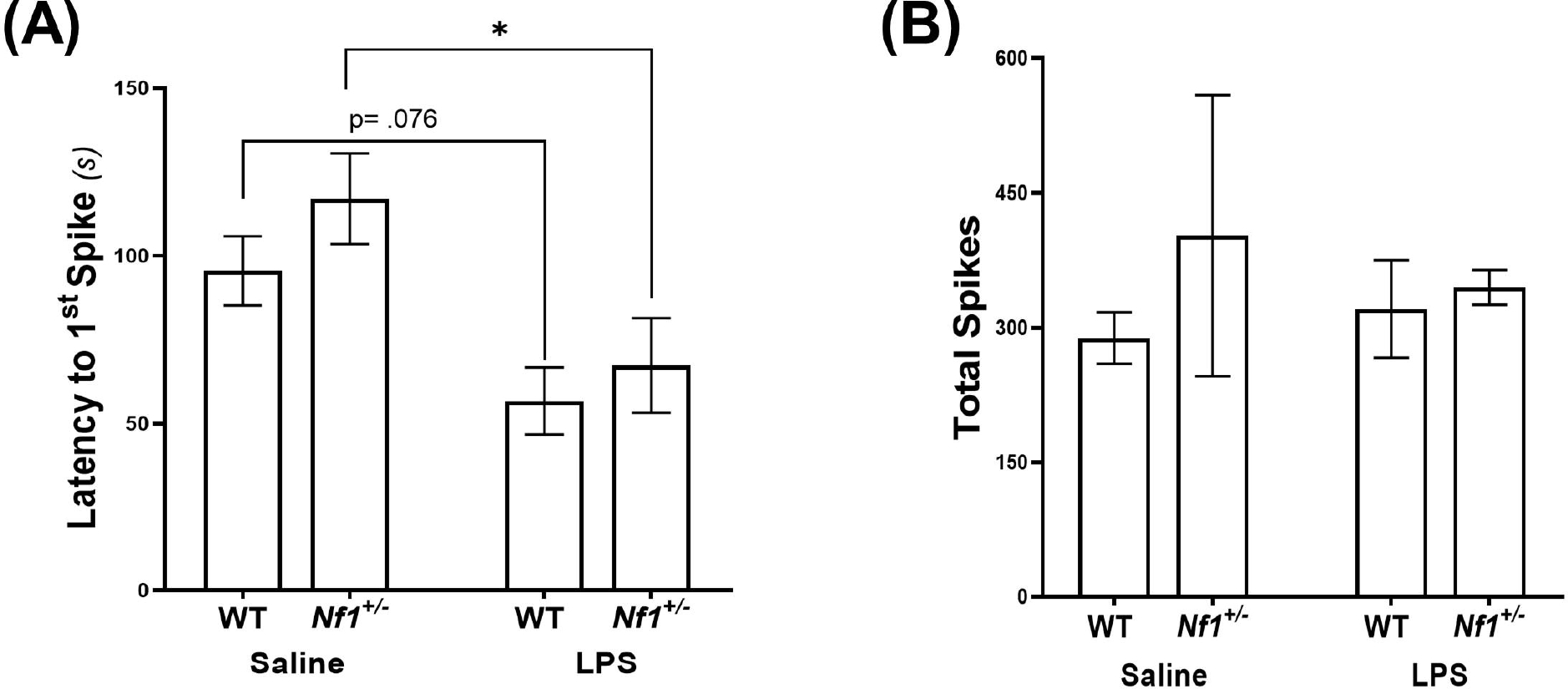
Early-life immune activation decreases the latency to the first spike, but does not affect the total spike count, following a PTZ challenge in both *Nf1*^*+/-*^ and WT adult mice. WT+LPS and Nf1+/-+LPS groups had significantly shorter latencies to the first spike after PTZ challenge as compared to the WT+Saline and Nf1+/-+Saline groups (A). There were no differences in the total spike counts in the hour following PTZ challenge (B). Data are presented as the mean ± SEM, *p<0.05.

The proportion of mice that developed seizures, the latency to the first PTZ-induced seizure and the cumulative seizure duration were also examined across groups. PTZ-induced seizures were detected in 2/8 WT+Saline mice, 2/6 *Nf1*^*+/-*^+Saline mice, 1/8 WT+LPS mice, and 2/6 *Nf1*^*+/-*^+LPS mice, with no statistical differences in the proportion of mice that developed PTZ-induced seizures (*χ*^*2*^ (3, *N*=28) = 1.11; *p*=0.774). There were no significant differences in the seizure latency or cumulative seizure duration (Appendix S2, Supplementary Figure 3).

## 4 Discussion

We set out to investigate whether early-life immune activation in NF1 further exacerbates the increased seizure susceptibility and learning and memory impairments associated with the disease. We found comparable basal neuroinflammation in P10 WT and *Nf1*^*+/-*^ saline-treated mice as well as a comparable cytokine neuroimmune response to peripheral LPS challenge. We have shown for the first time that adult *Nf1*^*+/-*^ mice have increased spontaneous epileptiform discharges compared to WT mice, as demonstrated in the saline control groups. We also demonstrate that early-life immune activation with LPS leads to a further increase in epileptiform abnormalities and promotes the development of spontaneous seizures in *Nf1*^*+/-*^ but not WT mice. These results support a role for early-life inflammation in the development of seizures and epilepsy in NF1. However, a similar effect of early-life immune activation was not found in spatial learning and memory tasks, suggesting inflammation may not be a factor in NF1-associated cognitive deficits.

The present study is the first to assess levels of pro-inflammatory cytokines in the brain of *Nf1*^*+/-*^ mice to determine the basal neuroimmune state of the CNS in NF1. There is evidence of increased concentrations of circulating inflammatory cytokines such as TGFβ (Farschtschi et al., 2016), interferon-γ, TNF-α, IL-6 (Farschtschi et al., 2016; Torres et al., 2016) and IL-1β (Park et al., 2013) in blood samples from patients with NF1. As such, we hypothesized that basal cytokine levels may be elevated in saline-treated *Nf1*^*+/-*^ *vs*. WT mice. We found, however, comparable baseline cytokine levels (TNF-α, IL-6, IL-1β), as well as a comparable cytokine neuroimmune response to LPS challenge. We cannot exclude, however, differences in other inflammatory cytokines and chemokines between *Nf1*^*+/-*^ and WT mice. We also cannot exclude any time-dependent differences in the brain cytokine response to the systemic LPS challenge, as has been previously described (Biesmans et al., 2013; Erickson and Banks, 2011).

This study is also the first to report spontaneous epileptiform abnormalities in *Nf1*^*+/-*^ mice. We found a significantly higher number of interictal spikes in saline-treated *Nf1*^*+/-*^ *vs*. WT mice, demonstrating for the first time that spontaneous interictal epileptiform activity occurs in adult *Nf1*^*+/-*^ mice. These results build on previous work from our laboratory in which we have shown an increased susceptibility to various chemical convulsants (Rizwan et al, 2019) and an increased rate of epileptogenesis (Sabetghadam et al, 2020) in adult *Nf1*^*+/-*^ mice. Although we did not detect any spontaneous seizures in *Nf1*^*+/-*^ mice that were not challenged with LPS, we cannot rule out the possibility that spontaneous seizure activity may occur at a low frequency which would require analysis beyond the five days of EEG recording analyzed in the present study. Both the frequency of interictal spikes and the occurrence of spontaneous seizures were increased in adult *Nf1*^*+/-*^+LPS, but not in WT+LPS mice. Although we have only reported on male mice, we do not have reason to believe there would be sex difference in terms of the effect of LPS on seizures and epilepsy in NF1. For one, our previous studies on seizure susceptibility and epileptogenesis in NF1 were performed on both males and females (Rizwan et al., 2019; Sabetghadam et al., 2020) with no sex differences reported. Similarly, sex differences have not been found in NF1 patients with epilepsy (Sorrentino et al., 2021). In addition, reports of increased seizures after early-life LPS in a mouse model of autism spectrum disorder also did not find sex differences (Lewis et al., 2018). Overall, our results support the hypothesis that the genetic mutation alone may not be sufficient to cause epilepsy in NF1, but that a second-hit from early-life inflammation may be a factor in promoting epileptogenesis.

We have previously reported an increased seizure susceptibility to pilocarpine and kainic acid in *Nf1*^*+/-*^ mice (Rizwan et al., 2019), and predicted they would also have an increased susceptibility to PTZ. We were therefore surprised to find no differences in PTZ-induced seizure parameters between WT and *Nf1*^*+/-*^ saline controls. The absence of an increased susceptibility of *Nf1*^*+/-*^ mice to PTZ, as opposed to the increased susceptibility to kainic acid and pilocarpine we have previously reported^*12*^, may be attributed to the differences in the mechanism of action of these chemical convulsants. PTZ is a non-competitive antagonist of the GABA_A_ receptor and elicits hyperexcitability by blocking GABA-mediated inhibition (Ramanjaneyulu et al., 1984), while kainic acid is an L-glutamate analog of the ionotropic kainate receptors (Nadler et al., 1978) and pilocarpine is a muscarinic acetylcholine receptor agonist (Turski et al., 1983). NF1 is associated with increased GABA tone in the hippocampus (Cui et al., 2008), which might negate some of the effects of PTZ and may explain why we did not find an increased susceptibility to PTZ in the *Nf1*^*+/-*^ mouse as we previously reported with other chemical convulsants (Rizwan et al., 2019).

Early-life systemic inflammation models have been shown to promote an increased seizure susceptibility to chemical convulsants in adulthood (Galic et al., 2008; Chen et al., 2013; Gomez et al., 2021). In the present study, early-life immune activation decreased the latency to first PTZ-induced spike in both WT and *Nf1*^*+/-*^ mice, but did not affect PTZ-related additional parameters as previously reported by other groups. This may be related to different techniques in assessing PTZ seizure susceptibility, such as the use of continuous tail vein PTZ infusion (Lewis et al., 2018; Gomez et al., 2021) versus the single systemic injection used here. However, for reasons mentioned above, PTZ may not be the most appropriate chemoconvulsant to use in NF1 studies going forward.

The mechanisms that underly NF1-associated seizure susceptibility have not been previously studied. Previous reports describe changes in voltage-gated ion channels associated with NF1 that may mediate hyperexcitability (Wang et al., 2010; Moutal et al., 2017). Furthermore, in addition to its interaction with Ras, neurofibromin is known to interact with NMDARs. While alterations in NMDAR have not been studied in the context of hyperexcitability in NF1, NMDAR activation has been shown to promote Ras-inactivation by neurofibromin and facilitate postsynaptic spine density and synaptic plasticity in pyramidal CA1 neurons (Husi et al., 2000). Enhanced ERK signaling has also been reported in the *Nf1*^*+/-*^ mouse model (Cui et al., 2008) and enhanced ERK signaling has been previously shown to stimulate expression of NMDA receptors (Li et al., 2005). If NMDAR trafficking is increased in NF1, then it is possible that an excitation/inhibition imbalance occurs, favoring hyperexcitability. The increased GABA tone previously described in NF1 may also promote hyperexcitability if limited to inhibitory neuronal circuits. In addition to these mechanisms, it is possible that early-life immune activation can promote epileptogenesis in NF1 by perturbing the developmental trajectory of the brain. In support of this hypothesis, previous reports have demonstrated that while LPS-stimulated *in vitro* cultures of *Nf1*^*+/-*^ microglia secrete comparable levels of pro- and anti-inflammatory cytokines as WT microglia, the phagocytic activity and purinergic signaling in adult *Nf1*^*+/-*^ microglia is significantly reduced (Elmadany et al., 2020). As microglia undergo morphological and functional changes throughout their developmental maturation (Schwarz et al., 2012), and given the role these cells play in fine tuning synaptic connections during neurodevelopment, it is possible for early-life immune activation to perturb the trajectory and function of these cells and affect the neuronal network in the NF1 brain.

*Nf1*^*+/-*^ mice were first described to have a spatial memory impairment in the Morris water maze (MWM) by Silva and colleagues, who demonstrated that the impairment is mild and can be overcome with additional acquisition training (Silva et al., 1997). Our data, the first describing the spatial learning and memory phenotype of *Nf1*^*+/-*^ mice in the Barnes maze, did not demonstrate a spatial memory impairment in adult *Nf1*^*+/-*^ mice. While we applied a common Bares Maze training regimen to what has previously been used in C57BL/6 mice (Sunyer et al., 2007), it is possible that the length of the acquisition phase promoted an overtraining of *Nf1*^*+/-*^ mice. Additionally, Silva and colleagues have reported that the impairment in spatial reference memory in *Nf1*^*+/-*^ mice is not fully penetrant and up to 30% of mice in their study had normal performance in the MWM. Taken together, while our data does not demonstrate a spatial memory impairment in the *Nf1*^*+/-*^ mouse, our results serve to guide future studies for phenotyping the *Nf1*^*+/-*^ mouse model in the Barnes maze. Early-life LPS challenge at P10 did increase the total distance travelled and the total number of errors in reaching the target hole in the probe trial in WT+LPS mice compared to WT+Saline mice, in contrast to the lack of effect of an early-life LPS challenge at P14 (Dinel et al., 2014). It is unclear why there was an effect of early-life inflammation in WT and not *Nf1*^*+/-*^ mice.

This is the first report describing the neuroinflammatory state and the long-term consequences of early-life immune activation in NF1. Our results have demonstrated that whereas a systemic early-life LPS challenge induces a *comparable* acute cytokine neuroimmune response in *Nf1*^*+/*-^ and WT mice, it promotes increased seizure activity only in adult *Nf1*^*+/*-^ mice. While *Nf1*^*+/-*^ heterozygosity increases epileptiform abnormalities, our findings show that *second-hit* environmental events (such as early-life immune activation) may be required to promote epileptogenesis. This may explain in part why some patients with NF1 develop epilepsy over their lifetime while others do not.

## Supporting information

Supplementary Results

## 7 Acknowledgements

We would like to thank Dr. Chiping Wu for his assistance with electrode implantation, and Dr. Mingdong Yong for assistance with maintaining the breeding colonies.

