## Supplementary Methods for "Early-life immune activation is a vulnerability factor for adult epileptogenesis in neurofibromatosis type 1"

*Supplementary Figures*

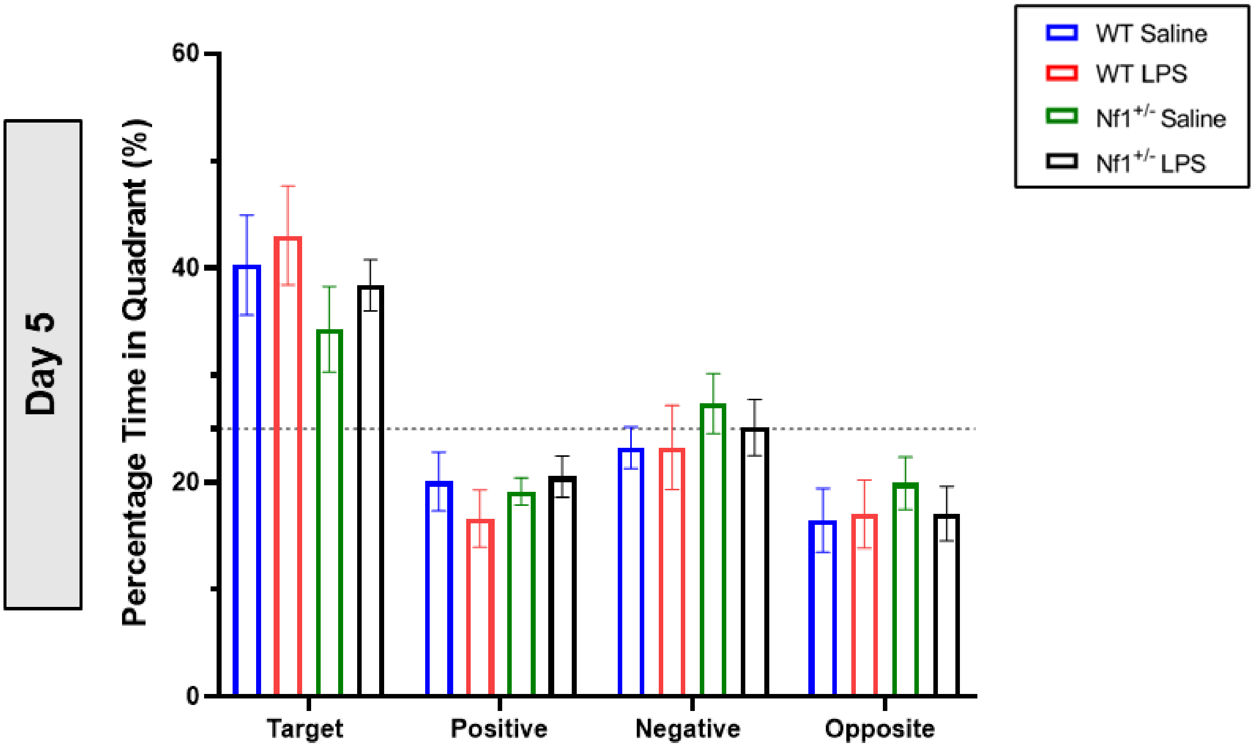


**Supplementary Figure 1. Quadrant Exploration time in the Barnes Maze.** Bar graphs represent the average time spent in each of the four quadrants (as a percentage of the total time) during the probe trial on Day 5 of testing. The target quadrant contains the target hole. The dotted line represents chance. Data are reported as mean ± SEM. Two-Way repeated measures ANOVA followed by Dunnett's post-hoc for within-group differences, Tukey's post-hoc for between-group differences. Control WT and *Nf1*$^{+/-}$ mice exhibited target quadrant preference during the probe trial: Control WT mice explored the Target quadrant significantly more than any other quadrant in the maze [Dunnett's post hoc (*vs.* Target), Positive: $p < 0.001$; Negative: $p = 0.002$; Opposite: $p < 0.001$] and control *Nf1*$^{+/-}$ mice showed bias towards the Target quadrant as well [(*vs.* Target), Positive: $p = 0.004$; Opposite: $p = 0.007$] although this was not generalized to all non-target quadrants [(*vs.* Target), Negative: $p = 0.306$]. However, there were no differences in the percentage of time spent exploring the Target quadrant when comparing between control WT and *Nf1*$^{+/-}$ mice. Similarly, both LPS-challenged WT and LPS-challenged *Nf1*$^{+/-}$ mice displayed target quadrant discrimination, as evidenced by exploring the Target quadrant significantly more than any other quadrant [WT (*vs.* Target), Positive: $p < 0.001$; Negative: $p = 0.001$; Opposite: $p < 0.001$, *Nf1*$^{+/-}$ (*vs.* Target), Positive: $p = 0.004$; Negative: $p = 0.041$; Opposite: $p < 0.001$]. The percentage of time spent exploring the Target quadrant was also not statistically different when comparing LPS-challenged WT and LPS-challenged *Nf1*$^{+/-}$ mice, and similarly not statistically different when compared to their saline-matched controls.

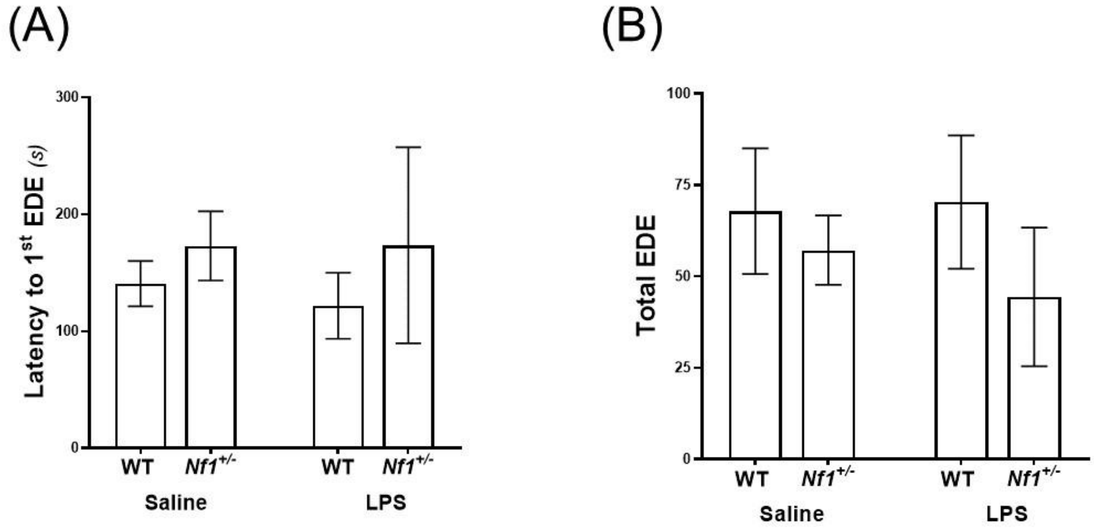


**Supplementary Figure 2. The latency to the first epileptiform discharge event (EDE) and total EDE count in *Nf1*$^{+/-}$ and WT adult mice exposed to LPS or saline as neonates**. Bar graphs represent the Mean ± SEM, and data were analyzed in a One-Way ANOVA. There were no statistical differences found in the latency to the first EDE nor in the total EDE count across all the groups.

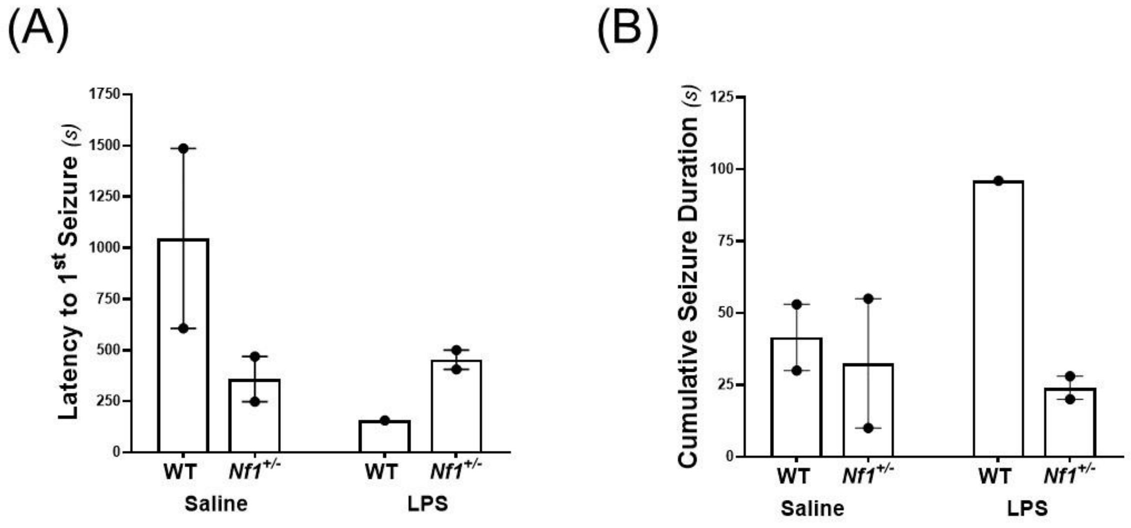


**Supplementary Figure 3. The latency to the first PTZ-induced seizure and cumulative seizure duration in *Nf1*$^{+/-}$ and WT adult mice exposed to LPS or saline as neonates**. An individual data point represents one mouse and, whenever appropriate, bar graphs displayed the mean ± SEM. PTZ-induced seizures were detected 3/8 control WTs, 2/6 control *Nf1*$^{+/-}$ mice, 1/8 LPS-challenged WTs and 3/6 LPS-challenged *Nf1*$^{+/-}$ mice with no statistical differences in the proportion of mice that developed PTZ-induced seizures ($\chi 2$ (3, N = 28) = 1.11; p = 0.774). The average latency to the first PTZ-induced seizure in control *Nf1*$^{+/-}$ mice appears to be shorter than that observed in control WT mice. The cumulative seizure durations, however, appear comparable across these control groups. Compared to their saline controls, LPS-challenged *Nf1*$^{+/-}$ mice appeared to have a comparable seizure latency and cumulative seizure duration. Only one LPS -challenged WT mouse developed PTZ-induced seizures and compared to the saline controls, appeared to have a shorter seizure latency and a longer cumulative seizure duration. However, statistical comparisons of these seizure latencies and cumulative seizure durations were not possible due to the small number of mice that developed PTZ-seizures.
