## Supplementary Results for "Early-life immune activation is a vulnerability factor for adult epileptogenesis in neurofibromatosis type 1"

### *Supplementary Methods*

#### *Brain tissue homogenization*

Brain tissue was thawed on ice and homogenized using a cordless motor pellet mixer (#V8185-904; VWR, Mississauga, Canada), in a PBS-based cell lysis buffer (#895347; R&DSYSTEMS, Oakville, Canada) complete with EDTA-free protease inhibitors (#04693159001;Roche, Mississauga, Canada) and 1 mmol/L phenylmethylsulfonyl fluoride (#PMS123.5;BioShop, Toronto, Canada). Tissue lysates were then spun in a centrifuge at 18,000 rpm for 20 minutes. The supernatant fraction was collected, and the pellet fraction was stored at -80C. Total protein was quantified using a detergent compatible assay (#5000113; BioRad, Mississauga, Canada). The concentration of protein in all samples was normalized to the same concentration using a calibrating buffer (#895438; R&DSYSTEMS, Oakville, Canada) as the diluent. All protein samples were stored at -80C until assayed in the Mouse Magnetic Luminex Screening Assay.

#### *Pro-inflammatory Cytokine Analysis*

Tissue lysates and standards were loaded in duplicates and analyzed according to kit standard instructions. Briefly, undiluted brain tissue lysates were incubated with a cocktail of magnetic microsphere beads, each pre-coated with analyte-specific antibodies and embedded with fluorophores at ratios set for unique bead regions. Following incubation, a wash step was performed to remove any unbound standard/sample and a cocktail of biotinylated detection antibodies, specific for the analytes of interest, was added to each well. Following incubation, and a round of washing to remove any unbound biotinylated antibodies, streptavidin-phycoerythrin conjugate (streptavidin-PE) was added. A final wash step was conducted to remove any unbound streptavidin-PE, the microparticles were resuspended, and the plate was analyzed in a Luminex® 200 analyzer. A classification laser (red, 635 nm) inside the analyzer excited the dye inside each bead to determine which analyte was being detected through the unique bead regions. A second laser known as the reporter laser (green, 532 nm) excited the PE reporter dye to measure the amount of analyte bound to each bead. Inside the analyzer, fluorescence emissions from each bead passed through a flow cell and were analyzed to differentiate their emission levels using a photomultiplier tube and an avalanche photodiode. Data for standards and samples from the Luminex® 200 analyzer were reported as median fluorescence intensity values, and a standard curve was generated for each analyte. Analyte concentrations (pg/mL) for each sample were extrapolated from the standard curves using a weighted five parameter regression analysis. These concentrations were normalized to the total protein value of every sample, then averaged and the mean analyte concentrations (pg/mg) were reported per experimental group.

***Barnes Maze***

Immune-challenged and (saline) control male and female mice were tested in the Barnes Maze to assess spatial learning and memory, as previously described (Sunyer et al., 2007). On day 1, mice were habituated to the paradigm by placing them in the middle of the platform under a cylindrical black chamber for 30 s, then allowed to explore the maze for 180 s to locate and enter the escape chamber. During habituation, the escape chamber was placed at a different hole than the target hole. If a mouse failed to find and/or enter the escape chamber within 180 s, it was gently guided to it. Once in the escape chamber, it remained inside for 60 s (by closing the chamber) and then was returned to its home cage. Following habituation, mice underwent a spatial acquisition phase spread across 4 days. During spatial acquisition, each mouse was trained to locate the escape chamber at the position of the fixed target hole over a 180s trial for 4 trials per day (with a 30-minute intertrial interval). Because some mice lacked motivation to enter the target hole and instead continued to explore the maze over time, parameters to the first encounter of the target hole (such as primary latency, primary distance and primary errors) as well as the mean speed were acquired and analyzed during the acquisition phase (Harrison et al., 2006). The primary latency and the primary distance were defined as the time (s) and the distance (cm) taken to the first encounter to the target hole, respectively. The number of head pokes into non-target holes before poking the target hole defined the errors measure. The first probe trial was conducted on day 5, 24 hours following the last training trial, to test short-term reference memory. The escape chamber was removed, and the animal was given 90 s to find the target hole. On day 12, a second probe trial was conducted, and long-term reference memory was similarly assessed. No tests were performed between day 5 to day 12. During each probe phase, the latency (s), distance (cm) and number of errors to the first encounter of the target hole, as well the mean speed (cm/s) and the percentage time spent in each quadrant of the maze were acquired. The number of errors was a measure that was scored manually by the experimenter during the acquisition and probe phase and multiple head pokes into the same non-target hole were counted as one error.
